# Positional isomers of a non-nucleoside substrate differentially affect myosin function

**DOI:** 10.1101/2019.12.17.879809

**Authors:** M. Woodward, E. Ostrander, S.P. Jeong, X. Liu, B. Scott, M. Unger, J. Chen, D. Venkataraman, E.P. Debold

**Affiliations:** University of Massachusetts at Amherst

**Keywords:** Molecular motors, muscle, control, laser trap

## Abstract

Molecular motors have evolved to transduce chemical energy from adenosine triphosphate into mechanical work to drive essential cellular processes, from muscle contraction to vesicular transport. Dysfunction of these motors is a root cause of many pathologies necessitating the need for intrinsic control over molecular motor function. Herein, we demonstrate that positional isomerism can be used as a simple and powerful tool to control the molecular motor of muscle, myosin. Using three isomers of a synthetic non-nucleoside triphosphate we demonstrate that myosin’s force and motion generating capacity can be dramatically altered at both the ensemble and single molecule levels. By correlating our experimental results with computation, we show that each isomer exerts intrinsic control by affecting distinct steps in myosin’s mechano-chemical cycle. Our studies demonstrate that subtle variations in the structure of an abiotic energy source can be used to control the force and motility of myosin without altering myosin’s structure.

**Statement of Significance:** Molecular motors transduce chemical energy from ATP into the mechanical work inside a cell, powering everything from muscle contraction to vesicular transport. While ATP is the preferred source of energy, there is growing interest in developing alternative sources of energy to gain control over molecular motors. We synthesized a series of synthetic compounds to serve as alternative energy sources for muscle myosin. Myosin was able to use this energy source to generate force and velocity. And by using different isomers of this compound we were able to modulate, and even inhibit, the activity of myosin. This suggests that changing the isomer of the substrate could provide a simple, yet powerful, approach to gain control over molecular motor function.

## Introduction

Molecular motors, such as muscle myosin, have evolved to transduce chemical energy from the gamma phosphate bond in adenosine triphosphate (ATP) into mechanical work. This energy is used to power a myriad of intracellular processes, from muscle contraction(1) to the transport of organelles(2), to cytokinesis(3). While ATP is the preferred source of molecular motors, there is growing interest in developing alternative energy sources to control biological motors and thereby control the processes they drive inside the cell(4, 5). For example, the ability to preferentially inhibit the activity of skeletal muscle myosin could aid in the treatment of muscle spasticity that is so prevalent in conditions like cerebral palsy. It would also be advantageous to activate or enhance skeletal muscle myosin function in disorders where neural input to the muscle is compromised, such as multiple sclerosis, ALS, and the “easy muscle fatigue” experienced by individuals with chronic heart failure (CHF). Directly enhancing cardiac myosin function would also be a powerful approach to improve the cardiac contractility in individuals with CHF. In fact, this approach is currently being pursued using 2-deoxy ATP, an analog of ATP (6–8). Thus, gaining control over molecular motors represents a promising area for novel therapeutic approaches to treat many prevalent conditions. Gaining control over molecular motors is also a focus of nanotechnology efforts that aim to build synthetic controllable molecular motor-based nanomachines(9–13). Indeed, biological motors have even been used, in this context, to perform parallel computing tasks(14), where precise control over the motors is critical to proper function of the nanodevice. Thus, there is growing interest in the ability to control molecular motor function both in vivo and in vitro because of the broad and powerful potential that such control would offer to both clinicians and bioengineers.

Several different strategies have been employed to gain control over the activation, speed, and directionality of molecular motors. Initial efforts using “caged-ATP”, enabled researchers to precisely control the initiation of contraction in muscle, providing novel mechanistic insight into the timing and sequence of the initial events of contraction. However, the degree of control with these types of compounds was limited to modulating the binding of the substrate to myosin’s active site, creating an irreversible “on/off switch”. Greater control can be gained by manipulating the surface chemistry of the experimental chamber in in vitro motility assays with isolated proteins. This strategy can be used to directly control both the pattern and speed of the filament motion driven by molecular motors(15). A related approach is to apply an external load to the charged actin filament by imposing a weak electrical field to the bathing solution in an in vitro motility assay(16). Using this approach, the velocity of the actin filament can be accelerated or decelerated by taking advantage of the load-dependent nature of myosin function(17, 18).

Alternatively, the structure of the motor can be altered using molecular biology techniques to introduce new structural elements to exert control. Using this approach, the structure of myosin’s motor domain can be altered such that its preferred direction of motion can be permanently reversed(19). Similar approaches have been used to incorporate photosensitive structural elements into the motor domain of myosin and kinesin that enable direct control over activation and direction of travel, which can be gated by light, with either UV(20) or blue light(11). While these approaches demonstrate an effective means of gaining control over motor function, they are more applicable to the development of in vitro nanodevices than to a clinical setting because the structural manipulations are too severe to be successfully implemented *in vivo*.

A better approach to control molecular motors in an in vivo setting is to custom design an alternative energy source (i.e. other than ATP) so that the activity of molecular motor could be controlled independent of its biological activation process. With this goal in mind, Menezes et al.(5) synthesized a triphosphate moiety onto the photo-switchable molecule, azobenzene. Using this compound myosin’s velocity could be regulated with exposure to different wavelengths of light in an in vitro motility assay(5). The bulky *cis* form of AzoTP (which predominates in UV light) likely renders AzoTP unable to bind to myosin’s nucleotide binding site, while the flatter *trans* form (that predominates with visible light exposure), readily binds to the active site much like ATP(5). Thus, this approach creates a type of control that is similar to that of the light-sensitive caged-ATP compounds(21), an “on/off switch”, but has the additional advantage of being reversible. However, the control over contraction is less precise than caged-ATP because conversion from the cis to trans form of AzoTP is incomplete; indeed even with prolonged exposure to UV 10% of the AzoTP remains in the *trans* form. This means that myosin function cannot be fully stopped. A further drawback for in vivo applications is that it would be difficult to implement an external light trigger in subcutaneous body tissues such as skeletal and cardiac muscle. Therefore, there is now a need to gain more precise intrinsic control over molecular motors, in which the substrate is modified to alter specific transitions within the molecular motor’s mechanochemical cycle (i.e. the cross-bridge cycle in myosin). This would provide a substantially greater degree of control over the motor such that both the activation and the speed of the motor could be precisely modulated rather than simply turning it on and off by affecting binding to the active site.

Gaining intrinsic control over molecular motors using alternative substrates requires a detailed mechanistic understanding of how myosin uses these compounds to generate force and motion. For example, despite the fact that substrates can be used by myosin to generate motion, it is not clear how they bind to and interact with myosin’s active site. It is also unclear which step or steps in myosin’s mechanochemical cycle might be altered to gain control over its function. Achieving this level of knowledge requires a full biophysical characterization of myosin’s mechanics and kinetics at both the ensemble and single molecule levels.

Herein we report that myosin can use a synthetic azobenzene-based non-nucleoside triphosphate (AzoTP) as an alternative energy source to generate force, as well as velocity, in in vitro assays. We show that myosin’s function is profoundly sensitive to the position of the triphosphate moiety (i.e. positional isomers) on the base of the synthetic substrate. In fact, with one isomer myosin was able to generate ~65% of the function elicited when ATP is the energy source, while another isomer reduced myosin’s velocity by ~99%; a third isomer inhibited myosin function in the presence of ATP in a dose-dependent manner. We used a series of biophysical experiments, including single molecule laser trap assays, to reveal how myosin used each of these AzoTP isomers to transduce energy, and compared the results with those using ATP. The results demonstrate that subtle structural changes to a synthetic substrate can dramatically alter and thus control myosin’s function by affecting the intrinsic properties of its mechanochemical cycle. Thus, positional isomerism provides a powerful new approach to gain control over molecular motor function and may also provide novel mechanistic insights into the fundamental nature, and efficiency, of energy transduction by this prototypical molecular motor.

## Materials and Methods

### Chemical synthesis

AzoTP compounds (Fig.1a) were synthesized based on a protocol reported previously(5). Briefly, we added di-tert-butyl *N,N*-diisopropylphosphoramidite, followed by oxidation with *m*-chloroperoxybenzoic acid to the ortho, meta or para isomers of hydroxy-2-ethoxy azobenzene in dry THF to form the *tert*-butyl protected monophosphate ester. The ester was deprotected with trifluoroacetic acid and was purified by eluting through a DEAE A-25 anion exchange column. During this process the cation was converted to a triethylammonium salt. A solution of triethylammonium monophosphate salt of the azobenzene was converted to a tributyl ammonium salt by the addition of tributylamine. This monophosphate was coupled with tributylammonium pyrophosphate to obtain tributylammonium salt of azotriphosphate. This was converted to its sodium salt using sodium iodide. The synthetic procedure of each compound is provided in the Supplemental Materials (Supplementary Fig. S1).

**Figure 1.**
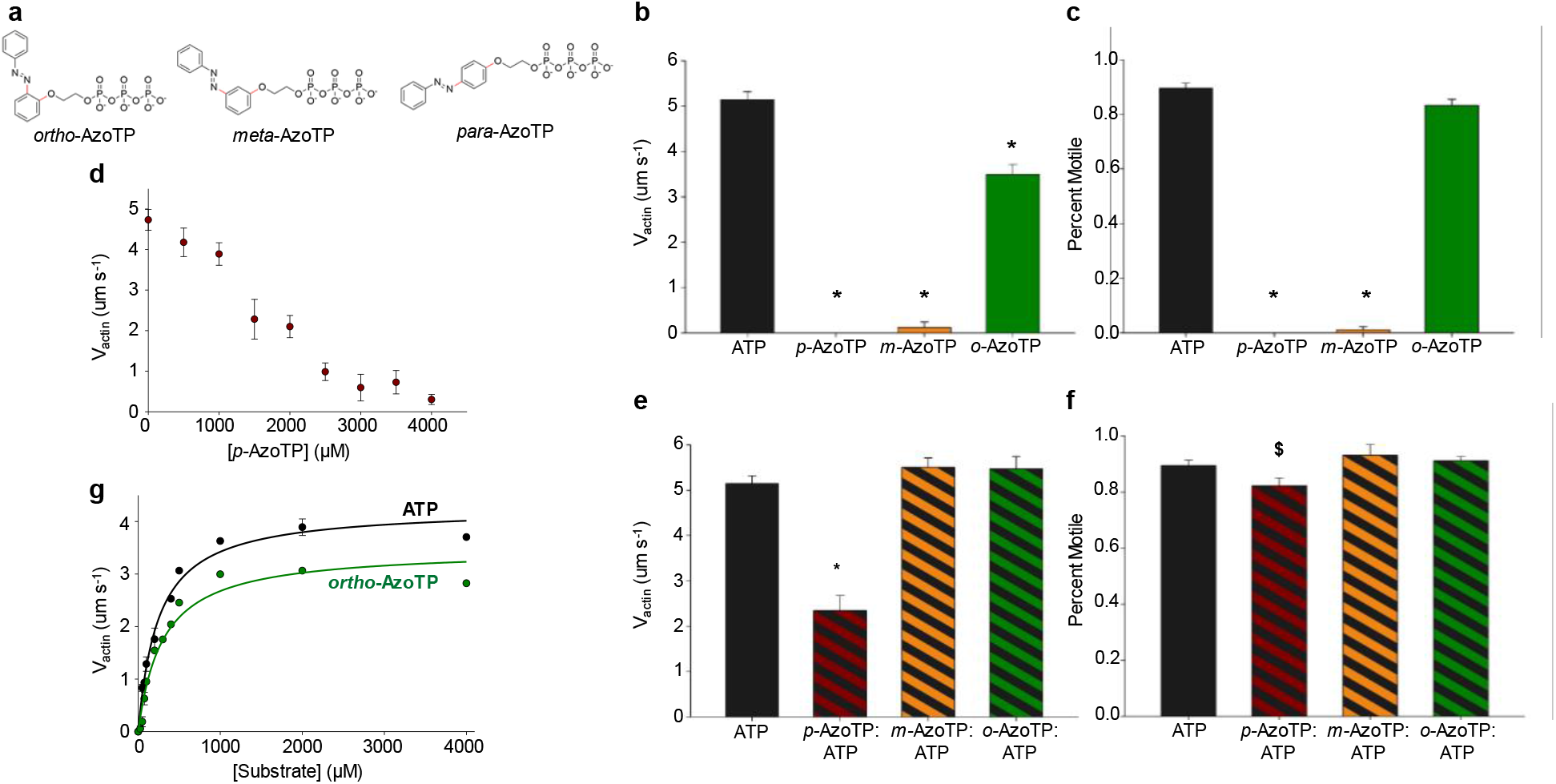
Effect of AzoTP isomers on myosin’s unloaded velocity. **a**) Chemical structures of isomers of AzoTP used in the functional assays as indicated. Note that the position of the triphosphate moiety is in the *ortho*, *meta* or *para* positions on the phenyl ring in each of the three structures. **b**) Bar graph showing the mean ± SEM of the actin filament velocities (V_actin_) of each substrate, including a control with ATP, at a concentration of 2mM. ATP, black (n = 17 observations); *para*-AzoTP red (n = 4); *meta*-AzoTP orange (n = 5); *ortho*-AzoTP green (n = 11). *indicates significantly (p<0.05) different from ATP. **c**) Mean ± SEM for percentage of actin filaments moving in **a**. **d**) Mean ± SEM V_actin_ values as a function of increasing concentrations of *para*-AzoTP in the present of constant (2mM) ATP (n = 3 - 5 observations at each concertation of para-AzoTP, except for 2000uM which was more, n = 14). **e**) Mean ± SEM V_actin_ values in the presence of mixtures of ATP and AzoTP isomers (2mM/2mM respectively) as indicated, solid black bar ATP control and stripped bars indicate V_actin_ values for compound mixtures (n = 5 - 8 for *para*-AzoTP/ATP, *meta*-AzoTP/ATP and *ortho*-AzoTP/ATP). **f**) Mean ± SEM for percentage of actin filaments moving in **e**, ^$^ indicates significantly different from *meta*-AzoTP value. **g**) V_actin_ as a function of substrate concentration. Mean ± SEM V_actin_ values for ATP (black dots) and *ortho*-AzoTP (green dots). Substrate concentrations ranged from 500uM to 4000uM. Each point represents 1-2 observations. Points fit to a Michaelis-Menten relationship, V_actin_ = V_max_*[S]/K_M_ + [S] yielding values for the maximum velocity (V_max_ = 4.3 ± 0.14 um·s^−1^ and 3.5 ± 0.22 um·s^−1^ for ATP and *ortho*-AzoTP respectively) and the apparent binding affinity of the substrate (K_M_ = 244 ± 25 uM and 298 ± 54 uM for ATP and *ortho*-AzoTP respectively). Both V_max_ and K_M_ were significantly (p<0.05) different between ATP and *ortho*-AzoTP.

### In vitro functional assays

#### Proteins

Myosin and actin used in the all the experiments were purified from chicken pectoralis muscle as previously described(22) with minor modifications as detailed previously(23). Roughly >95% of the myosin in this tissue is composed on the fast type IIb myosin heavy chain isoform(24). The action was isolated as G-actin but then re-polymerized and labeled with 100% TRITC/phalloidin for the in vitro motility assay, and with 50% TRITC/phalloidin and 50% biotin/phalloidin for the laser trap assay experiments. All animal tissue was obtained in accordance with the policies of the National Institute of Health using a protocol approved by the Institutional Animal Care and Use Committee at the University of Massachusetts Amherst.

### In vitro motility assay

The in vitro motility assay was performed as previously described(25), with minor modifications. Briefly, myosin was loaded onto a nitrocellulose-coated coverslip surface at a saturating concentration of 100 μg/mL. The surface was then blocked with BSA for 1 minute, and an unlabeled filamentous actin was added to prevent any rigor myosin molecules (aka dead-heads) from slowing the labelled actin velocity. TRITC-labeled actin was then added in the absence of ATP. The final buffer was added, containing either 2 mM ATP, 2 mM of an AzoTP compound, or a combination of 2 mM ATP and 2 mM of an AzoTP compound. Filament motion was visualized using a Nikon Ti-U inverted microscope, with a 100x, 1.4 NA CFI Plan Apo oil-coupled objective with the temperature maintained at 30.0 °C for all experiments. For each flow cell, three 30 s videos were captured at 10 frames/s and at three different locations within each flow cell.

### Analysis of in vitro motility data

The velocity of the actin filaments as well as the percent of filaments moving was determined using an automated filament-tracking ImageJ plugin WRMTRK. Filaments shorter than 0.5 μm were eliminated from the analysis, and filaments with velocities less than 0.13 μm·s^−1^ were considered to be stationary. A typical field of view generated 25–50 filament velocities, and the mean of these velocities was taken as the average velocity for that field of view, with a total of three recordings done for each flow cell. For each condition tested, at least three flow cells were used to generate the data, resulting in at least nine recordings contributing to the overall mean filament velocity for each condition.

### Single molecule and mini-ensemble laser trap assays

The laser trap assays were performed at 25 °C as previously described(26) and the system calibrated with established methods(27). Myosin was loaded into a flow-cell with number 1 thickness, nitrocellulose coated, coverglass (Thermo Fisher Scientific) in a high salt buffer (300 mM KCl, 25 mM imidazole, 1mM EGTA, 4 mM MgCl_2_, 1 mM dithiothreitol). For single molecule experiments myosin was loaded at 0.05 μg·mL^−1^, but for mini-ensemble experiments it was increased to 25 μg·mL^−1^. The surface was then blocked with BSA (0.5 mg/mL) before a low-salt experimental buffer (25 mM KCl, 25 mM imidazole, 1 mM EGTA, 4 mM MgCl_2_) with appropriate concentration of ATP or AzoTP. The experimental buffer also contained 1 μm diameter silica beads (Bangs Laboratories Inc.) coated with neutravidin, providing a linkage for attachment to the biotin/TRITC-labeled actin filaments. Two of these 1μm neutravidin-coated silica beads were trapped in time-shared laser traps and a single TRITC/biotin coated actin filament was attached to both beads in a three-bead configuration previously detailed(28). Compliance of this bead-actin-bead assembly was minimized by applying 3-4 pN of pretension, and the laser power set such that the combined trap stiffness of the dumbbell was 0.02 pN per nanometer, which was determined based on the Brownian motion of the bead-actin-bead assembly when not attached to myosin using equipartition theory(29). The dumbbell was then brought into contact with the myosin head(s) on the surface by approaching the nitrocellulose coated 3 μm silica bead.

### Analysis of laser trap assay data

To determine the size of myosin’s powerstroke (*d*) and the duration of actomyosin strong binding (*t*_on_) the raw actin displacement data were analyzed using a mean-variance analysis as previously described(30). Briefly, the mean and variance of the raw displacement records were calculated within a defined time window (e.g. 30 ms). This window was advanced over the entire displacement record and the calculated mean and variance within each window was plotted in a 3-D histogram with the third dimension indicating the number of observations binned within a specific mean variance value. Myosin’s step size was taken as the distance in nanometers from the center of the fit to the unbound lifetime (baseline, B) and the center of the event population (*e*).

Varying the window width over a single displacement record provides a measure of the density of binding events at each window width. Event density (*p*) as a function of the window width (*tw*) was then plotted and fit with a mono-exponential decay, with *t*_on_ the amount of time myosin spends strongly bound to actin and *N* the number of binding events, using the following equation:

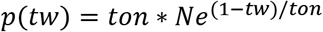

A separate custom analysis program was used to analyze the mini-ensemble laser trap assay data, to determine peak force and event duration for all recordings. This threshold based program was similar to the method we previously published(31) with minor modifications, including automated detection(26). Briefly, the data was processed with a custom Python script to correct for baseline wander. Analysis of the processed data, including the event detection and measurements, were completed in R^©^ 3.5.2(32) using custom programs with functions from the ‘*tidyverse*’ and ‘*gtools*’ packages(33). Briefly, the baseline corrected data was low-pass filtered using a running mean with a 10 ms (50 data points, at the sampling rate of 5 kHz) window width and the events were identified using two criteria, 1) an +8 nm threshold was set to the running mean to signal the start/end of an event, and 2) the event was included in further analysis if the duration was greater than 10 ms. Event durations were calculated as the time between the start/end (8 nm) thresholds and the time-off measurements were calculated as the time between the end of an event and the start of the subsequent event. Peak forces were calculated by identifying the maximum displacement of each record and converting the displacement in nanometers to forces in pN (0.04 pN·nm^−1^). Plots of the raw data and representative analysis traces were constructed using the ‘*ggplot2*’ and ‘*cowplot*’ R^©^-packages(33, 34). The data were then plotted in a histogram (Fig. 4b) and the arithmetic means reported (Fig. 4d-f).

### Molecular modeling and dynamics simulations

Atomistic simulations of myosin in complex with ATP or Azo-TP molecules were carried out using CHARMM/OpenMM program(35, 36). The latest CHARMM 36 m force field(37) was applied to model protein, water and ions, and CHARMM general force field (CGenFF)(38) was used to model the small molecules. The initial protein structure was obtained from RCSB Protein Data Bank (PDB) entry 1MMD(39). Coordinates of Mg^++^ and water molecules from this PDB structure were also retained. ATP and AzoTP molecules were first generated using CHARMM-GUI(40, 41). The ligand was then translated to the binding pocket of myosin, with the coordinates of TP group replaced by those of adenosine diphosphate•BeF_3_ in PDB 1MMD and the Azo group reoriented to mimic the binding pose of ADP. The initial structure of myosin-ligand complex was energy minimized in vac
uum for 5000 steps to remove any steric clashes. It was then placed in a rectangular simulation box containing ~28,000 TIP3P water molecules, such that the minimum distance between protein surface and box edge was at least 12 Å. 9 K^+^ ions were also added to neutralize the system. The simulation box was ~126 * 96 * 78 Å(37), and periodic boundary condition was imposed for all simulations.

The solvated systems were energy minimized using steepest descent and adopted basis Newton-Raphson algorithms. Each system was then slowly heated up from 100 K to 300 K in a 10-ps simulation, with the heavy atoms of protein and ligand restrained by harmonic potentials with a force constant of 5 kcal/mol/Å(36). Another 1 ns simulation was also performed to equilibrate the system at 300 K and 1 atm, with the retraining potentials slowly reduced to zero. The final production runs were carried out under the same NPT conditions without any restraint, except that the Azo group was restrained at the *trans*- conformation with a harmonic potential of force constant 5 kcal/mol/Å(36). In all simulations, the nonbonded interactions were truncated at 13 Å, and long-range electrostatic interactions were treated with particle mesh Ewald method(42). Lengths of all bonds involving hydrogen atoms were constrained using SHAKE algorithm(43). The time step to integrate the equation of motion was 2 fs, and final production runs lasted for at least 100 ns.

### Additional statistical analyses

Comparisons among the single molecule laser trap assay and motility data were determined using ANOVA and Tukey’s post-hoc tests (Sigmaplot 11.0) to locate differences with the alpha level set at p < 0.05. A Student’s t-test was used to determine if the ortho-AzoTP resulted in a significant difference in myosin’s step size, duration of strongly-bound lifetime (*t*_on_) vs ATP.

The nonlinear regression for estimating the xDP-release and xTP-binding rates (Fig. 5) were performed in *R*^©^ version 3.5.2 using the *nls* function from the *stats* package(32) which optimizes the fit using the least squares method (Gauss-Newton algorithm). Summary statistics from the regression were extracted using the *tidy* function from the *broom* package(44). These data were visualized using the *ggplot2* package(33). The goodness of fit for regression was determined through the residual plots using the *nlsResiduals* function in the “*nlstools*” package(45).

## Results

### Positional isomers of AzoTP differentially affect myosin function

We synthesized three positional isomers of an azobenzene-based triphosphate (AzoTP) to use as non-nucleoside energy sources for fast skeletal muscle myosin. The phosphate moiety was placed at the *ortho*, *meta* or *para* positions of one of the phenyl rings (Fig. 1). Each compound was tested to determine its ability to power muscle myosin as it translocated actin filaments in an in vitro motility assay (Fig. 1b). The velocity of the actin filaments in the motility assay (V_actin_) was determined and compared with the value observed when ATP served as the control substrate (Fig. 1b). When *para*-AzoTP was used as the energy source, myosin did not move the actin filaments (Supplemental video 1), suggesting it cannot harness the energy in the triphosphate. Likewise, no filament motion was initially observed when using *meta*-AzoTP; however upon closer examination over a more extended time scale (2min vs 30sec), there was clear evidence of actin filament motion with *meta*-AzoTP, albeit with an extremely slow velocity of 70 nm per second, less than 1% of the velocity observed with ATP (Supplemental video 2). Indeed, almost all of the filaments (~90%) moved at a constant velocity of 70 nm/s, suggesting that myosin is still using the energy of *meta*-AzoTP, but that one or more steps in its mechanochemical cycle was drastically slowed. In contrast to the *para* and *meta* isomers, with *ortho*-AzoTP serving as the energy source, myosin generated a V_actin_ that was ~65% of that observed using ATP (Fig. 1a and Supplemental video 3), and as with ATP, >80% of the filaments were motile (Fig. 1c). These findings demonstrate that myosin can harness the energy from a non-nucleoside triphosphate to power actin filament motion.

50/50 mixtures of each AzoTP variant with ATP were investigated to determine if the compounds could inhibit myosin function in the presence of ATP. When *meta* or *ortho*-AzoTP were mixed with equal amounts of ATP, V_actin_ was similar to ATP alone, suggesting that they do not affect myosin’s interaction with ATP and thus likely do not inhibit myosin function (Fig. 1d). In contrast, *para*-AzoTP slowed V_actin_ by 50% when it was mixed with an equal amount of ATP (2mM) (see Supplemental movie 4). The percentage of moving filaments in this mixture was equal to that of ATP alone (Fig. 1c), suggesting that the decreased velocity was not due to an increased portion of non-moving filaments.

To determine the nature of this inhibition we examined the effect of *para*-AzoTP on V_actin_ as a function of the concentration *para*-AzoTP, initially at a constant ATP concentration of 2mM (Fig. 1d). This revealed a linear decrease in V_actin_ as a function of the concentration of *para*-AzoTP. To further delineate the mechanism inhibition we measured the degree of inhibition of three different *para*-AzoTP concentrations while varying the ATP concentration. These findings indicated that the inhibitory effect of *para*-AzoTP did not increase as the ATP concentration was decreased (Supplementary Fig. 2), suggesting that *para*-AzoTP acts as an uncompetitive inhibitor of ATP(46). Importantly, the inhibitory effects were readily reversible, because washing out the *para*-AzoTP with a 2mM ATP buffer almost completely restored V_actin_ to the control (ATP) value (Supplemental video 5).

The ability of myosin to use *ortho*-Azo-TP to generate actin filament velocities in the motility assay at 65% of the value with ATP (Fig. 1b) enabled us to identify the steps in myosin’s mechanochemical cycle that might be altered when by the synthetic substrate. We started this effort by measuring V_actin_ as a function of the substrate concentration (Fig. 1g), and found that V_actin_ was slower with *ortho*-AzoTP at all substrate concentrations, with the maximum velocity (V_max_) observed to be 65% of that of the value with ATP. Notably at low substrate concentrations (<100uM) the filaments stopped moving with *ortho*-AzoTP, but filament motion was detected down to 50uM when ATP was the substrate. These data were fit to a Michaelis-Menten relationship, and indicated that the K_M_ for *ortho*-AzoTP was higher than for ATP, suggesting that *ortho*-AzoTP has a lower affinity for myosin’s nucleotide binding site. However, the decrease in V_actin_ at saturating substrate concentrations (>1mM) suggested that either myosin’s powerstroke is decreased and/or the rate of release of the diphosphate form of the substrate was slower than with ATP(47).

### Ortho-AzoTP affects myosin’s single molecule mechanics and kinetics

To more clearly delineate the mechanism underlying the decreased V_actin_ with *ortho*-AzoTP vs. ATP, we directly measured the size of myosin’s powerstroke (i.e. step size) and the duration of actomyosin binding (*t*_on_) using a single molecule laser trap assay (Fig. 2a).

**Figure 2.**
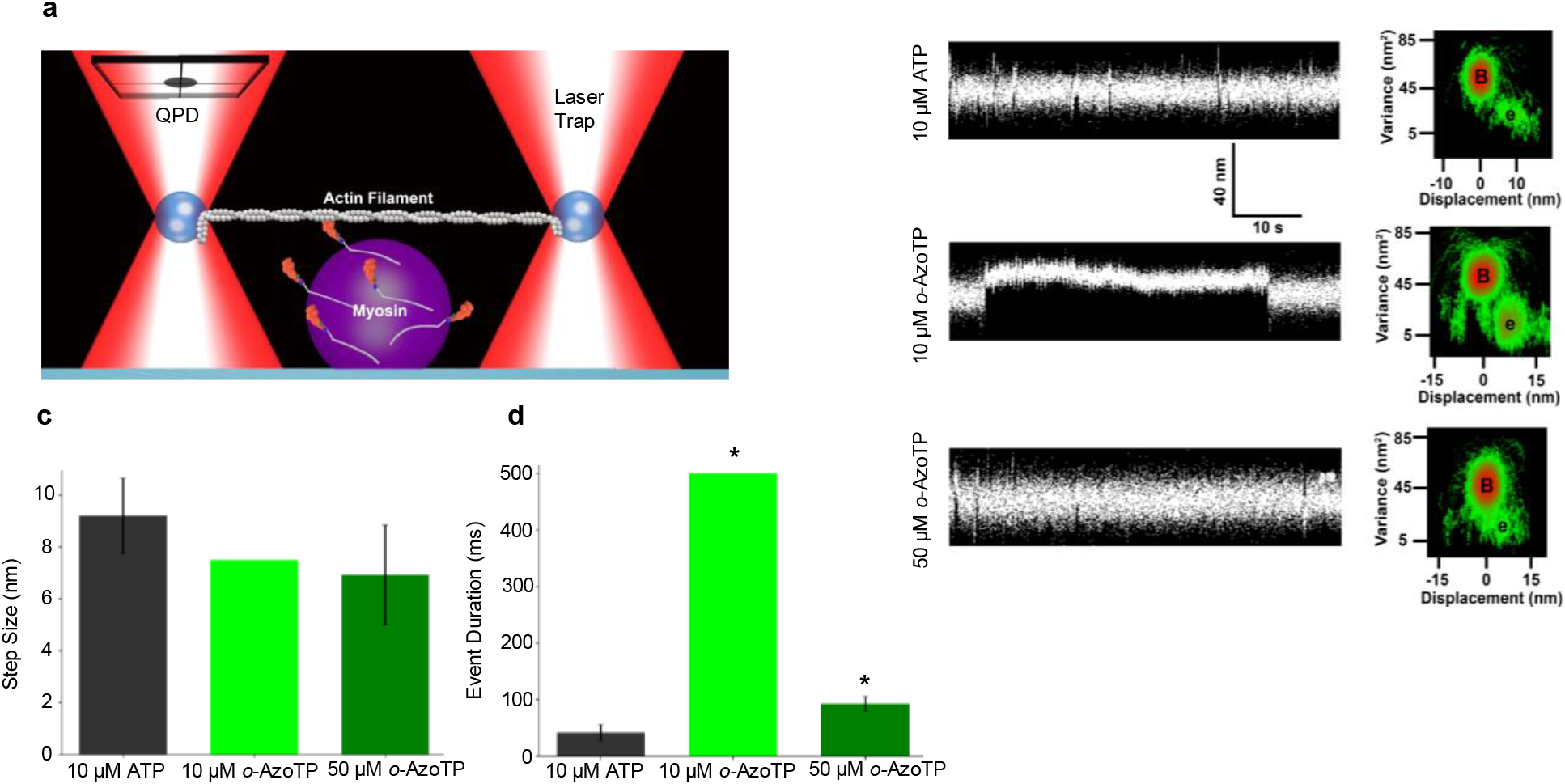
Effect of *ortho*-AzoTP on myosin’s step size and binding event duration. **a**) Schematic of the single molecule laser trap assay to determine myosin’s single molecule step size and binding event duration (see Methods). Actin filament displacement was tracked using a quadrant photodiode (QPD). **b**) Raw actin filament displacement records from the single molecule laser trap assay at 10 uM ATP and at 10 and 50 uM *ortho*-AzoTP. Actomyosin binding events are visible as reductions in the noise of the signal and a shift in the mean displacement. Binding events were much longer using *ortho*-AzoTP than ATP at 10uM ATP. These data were analyzed using the Mean-Variance analysis (see Methods) which calculates the mean and variance within a defined window of the data (e.g. 50ms). The window is moved over the entire record and the results plotted as mean displacement vs variance, with color intensity indicated the amount time spent at each value (red indicates highest intensity, green the lowest). The resultant 3-D histogram of the data reveals a baseline population (B) corresponding to the periods when myosin is not strongly-bound to actin, and event population (e) corresponding to periods of actomyosin strong-binding. The step size is determined as the difference between the B an e along the displacement axis. **c**) Mean ± SEM of myosin’s single molecule step size in nm. The total number of actomyosin binding events for 10uM ATP and 50uM *ortho*-AzoTP were (n= 202 and n= 67 respectively). The mean step size was determined from the MV-analysis of the data records was 9 ± 2 nm and 7 ± 2 nm for ATP and AzoTP respectively. Due to the extremely long binding event durations the total number of binding events for 10uM AzoTP (n = 56) was much less than for 10uM ATP (n = 202) despite a longer duration of recording time (375 and 260 seconds for AzoTP and ATP respectively). **d**) Mean ± SEM event durations at the indicated substrate concentrations. These were determined by plotting the event density as a function of window width using the Mean-variance analysis (see Methods).

Compared to ATP, myosin produced a step size (7 nm) that was 20% smaller with *ortho*-AzoTP (Figs. 2b & 2c), although not significantly different than that observed with ATP (9 nm). More striking, at 10 μM, the lifetime of a single actomyosin interaction (*t*_on_) was nearly 10-fold longer with *ortho*-AzoTP vs ATP (Fig. 2c). Indeed, even when the substrate concentration was increased to 50 μM the binding event lifetimes with ortho-AzoTP remained significantly longer than those at the 10 μM ATP (Fig. 2c). An equivalent measure of *t*_on_ at 50 μM ATP could not be obtained because binding events were shorter than the resolution of the instrument (~5-7 ms). The longer *t*_on_ observed in the single molecule laser trap assay is consistent with the lower K_M_, from the in vitro motility assay data (Fig. 1g). Indeed, the single molecule experiments provide much greater spatial and temporal resolution than in the bulk motility assay, where quantifying V_actin_ at μM substrate concentrations can be beyond the resolution of the instrumentation. Thus, these data suggest that *ortho*-AzoTP either binds much more slowly to myosin in the rigor state or that *ortho*-AzoTP is much slower to initiate the structural events that cause myosin’s dissociation from the actin filament(48).

We gained further insight into the mechanism of transduction by combining the step size (*d*) determined from the single molecule assay (Fig. 2), with the measurements of V_actin_ as a function of the substrate concentration in the motility assay (Fig. 1g), using a simplified model of myosin’s cross-bridge cycle (Fig. 3). This model estimates the rates of: a) the release of the diphosphate form of azobenzene triphosphate (*k*_−AzoDP_) from myosin; and b) the rate of AzoTP binding (k_+AzoTP_) to myosin’s active site(49). The model assumes that *t*_on_ is composed of the time waiting for the diphosphate form of the substrate to dissociate from myosin (inset Fig. 3), and the time waiting for the next triphosphate molecule to rebind to myosin and cause dissociation from actin. This was fit to the motility and single molecule data (Fig. 3) using the following equation:

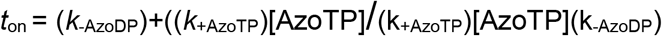

where *t*_on_ is the duration of strong actomyosin binding (obtained by dividing the single molecule step size, *d* (Fig. 2c) by V_actin_ at each substrate concentration (Fig. 3)); *k*_−AzoDP_ is the rate of release of the diphosphate form of AzoDP; *k*_+AzoTP_ the second-order binding constant of AzoTP to the rigor state; and [AzoTP] the substrate concentration.

**Figure 3.**
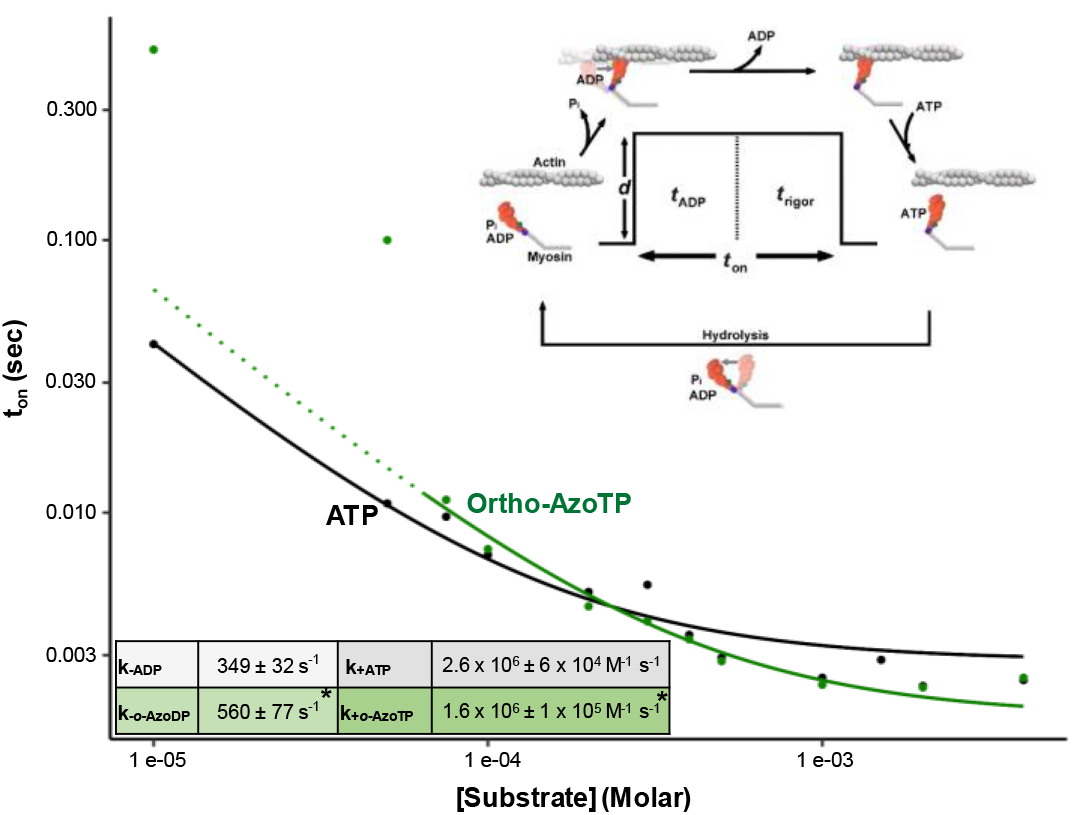
Effect of substrate concentration of myosin’s kinetics and mechanics. Main graph shows calculated or measured actomyosin event durations plotted as a function of substrate concentration for ATP (black) and *ortho*-AzoTP (green). Event durations (*t*_on_) from 4mM down to 75uM were determined by dividing the average single molecule step size (Fig. 2c) by V_actin_ at the corresponding substrate concentration (see Fig. 1). Event durations for 10uM ATP and 10 and 50uM AzoTP were taken from the values obtained in the single molecule laser trap assay (Fig. 2d). The data were fit to an equation (see Methods) based on a simple model of myosin’s cross-bridge cycle (inset figure). However, the fit could not converge to capture the long event durations in the single molecule assay observed at 10 and 50uM *ortho*-AzoTP. The dotted line indicates the extrapolated fit ignoring these two data points. The fit yielded values for the rate of the 2^nd^ order xTP induced binding constant (*k*_+ATP_ and *k*_+AzoTP_). The value for *k*_+AzoTP_ was significantly slower that for *k*_+ATP_, indicated by * in the inset Table. The fit also indicated that the rate of xDP release (*k*_−AzoDP_) was significantly faster for AzoTP.

The results of modelling indicated that *k*_+AzoTP_ was ~50% slower for *ortho*-AzoTP than for ATP (Fig. 3), consistent with a lower K_M_ from the Michaelis-Menten fit to the V_actin_ data (Fig. 1g). In contrast, the model suggested that myosin released AzoDP almost twice as fast as it released ADP (Fig. 3). These findings suggest that the slower V_actin_ is due to a slower rate of AzoTP binding to, and dissociating myosin from, the actomyosin rigor complex, combined with a reduced step size. Indeed, the substrate-induced dissociation was so much slower at 10 and 50 μM that these data points could not be fit by the model (Fig. 3), suggesting that k_+AzoTP_ is likely even slower than our model estimates. Interestingly, the reduction in V_actin_ comes despite a significant increase in the rate of release of the diphosphate form of the substrate AzoDP, adding further evidence that *k*_+AzoTP_ is much slower than with ATP.

### Myosin can harness the energy in ortho-AzoTP to generate force

To gain a more complete assessment of myosin’s ability to transduce energy from a synthetic energy source we also quantified myosin’s force generating capacity with *ortho*-AzoTP. This was done by using the laser trap assay as a picoNewton (pN) force transducer, where force is generated by a mini-ensemble of myosin molecules (Fig. 4a). Under the control condition with ATP serving as the substrate, the mini-ensemble of myosin molecules (~8) bound to and translocated the actin filament against the spring-like load of the laser trap (Fig. 4b). The stochastic nature of myosin binding and unbinding to the actin filament causes a distribution of forces to be generated, with peak forces ranging from 0.2 – 3.5 pN, and an average force equal to 1pN (Fig. 4c).

**Figure 4.**
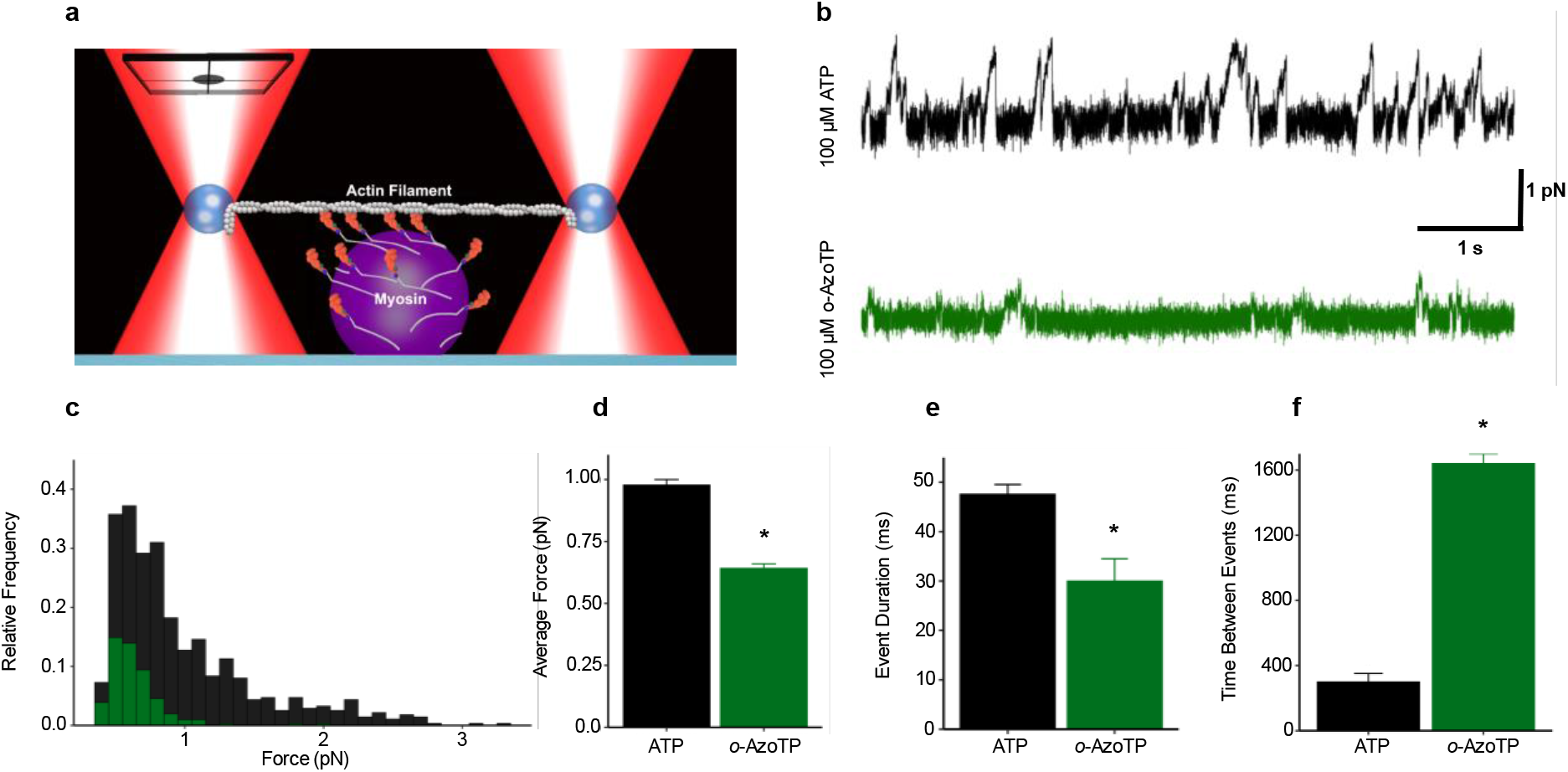
The effect of ortho-AzoTP on myosin’s force-generating capacity. **a**) A schematic of the mini-ensemble laser trap assay used to determine myosin’s force generating capacity. The assay is identical to the single molecule laser trap assay (Fig. 2a) except that the concentration of myosin of the surface is increased such that ~8 myosin heads are available to interact with the single actin filament and the substrate concentration was increased to 100uM. Force was determined from the displacement records by multiplying the displacement (*d*) by the trap stiffness (i.e. Force = *k*_*trap*_ * *d*). Trap stiffness was typically 0.02 pN·nm^−1^ as determined by the equipartition method (see Methods). **b**) Raw force records for ATP (black) and *ortho*-AzoTP (green). Multiple myosin molecules bind to the actin filament creating runs of motility against the increasing force of the laser trap. **c**) Peak force for each binding event determined using a custom algorithm (see Methods) was plotted in a histogram for each substrate. The total number of events for ATP 692 was and force *ortho*-AzoTP was 159. And the total amount of all data records was 274 sec. for ATP and 309 sec. min for *ortho*-AzoTP. **d**) Mean ± SEM peak force for each substrate. The mean forces were; 0.98 ± 0.02 pN for ATP and 0.64 ± 0.02 pN for *ortho*-AzoTP **e**) Binding event duration as determined by a custom algorithm (see Methods) and was 48 ± 2ms for ATP and 30 ± 4 ms for *ortho*-AzoTP. **f)** The time between binding events was also calculating using our custom algorithm and was 303 ± 23 ms for ATP and 1644 ± 198 ms for *ortho*-AzoTP. * indicates significantly (p<0.05) different from ATP.

In the presence of *ortho*-AzoTP myosin was able to generate roughly 65% of the force observed with ATP (Fig. 4b). Representative raw records show that the runs of motility of these ensembles were shorter than with ATP (Fig. 4b), despite the experiments being performed with the same number of myosin molecules and the same trap stiffness. A full analysis of all binding events revealed a reduction in the number and amplitude of force generating events (Fig. 4c), resulting in 35% less peak force than with ATP (Fig. 4d). Consistent with the reduction in force, the average lifetime of the binding events was reduced by roughly the same amount as the force (38%, Fig. 4d). This suggests that the drop in force, relative to ATP, is due to a slowed rate of attachment of myosin heads to the actin filament; the longer time between binding events adds further support for this hypothesis (Fig. 4e). Despite the reduced force levels, these data do demonstrate that this variant of a non-nucleoside synthetic triphosphate compound can be used by myosin to generate force over a distance, and thus mechanical work, in addition to its ability to propel actin filaments (Fig. 1b).

Similar force measurements could not be made using para-AzoTP because myosin did not generate any movement when it was used as a substrate (Fig. 1b). Likewise, the extremely slow velocity generated when *meta*-AzoTP was the substrate (~1% of that with ATP), prohibited the ability to accurately measure the force generating capacity with this substrate.

## Discussion

We successfully demonstrated that myosin can transduce chemical energy from a photo-switchable, non-nucleotide triphosphate compound into mechanical work (Figs. 1 & 4). Furthermore, by changing only the position of the triphosphate moiety from the *ortho* to *para* positions, AzoTP can be transformed from an energy source into an inhibitor of myosin function in the presence of its preferred energy source ATP. In addition, by moving the triphosphate to moiety to the *meta* position, myosin function can be slowed to <1% of the ATP-driven velocity. Thus, we have demonstrated that myosin’s force and motion generating capacity can be intrinsically controlled using a non-nucleoside-based molecule, enabling us to “shift the gears” of this molecular motor. Furthermore, by manipulating the substrate concentration and changing the number of myosin molecules in the assays, we have gained insight into the mechanisms underlying this control over the motor’s “transmission”.

Previous reports of using light-sensitive compounds to power muscle myosin found that specific variants of azobenzene-triphosphate can generate actin filament velocities up to 70% of ATP(5, 50). However, only molecules with the triphosphate moiety in the *ortho* position were examined, and only under unloaded conditions (e.g. in vitro motility assay or ATPase), in which no external work is performed by the motor. Therefore, our use of the mini-ensemble laser trap assay demonstrates, for the first time, that myosin can harness the energy from AzoTP to produce force against a load and therefore generate mechanical work using a substrate other than its preferred source, ATP (Fig. 4). This finding provides novel insight into the mechanism of myosin’s energy transduction because myosin’s force generating capacity is thought to be limited by distinctly different mechanisms than its unloaded velocity(47, 51, 52). In addition, our systematic manipulation of the position of the triphosphate moiety on the azobenzene molecule offers detailed insight into how the substrate might interact with myosin’s nucleotide binding site.

### Ortho-AzoTP slows actomyosin dissociation but accelerates AzoDP release

While myosin was able to generate force and velocity using *ortho*-AzoTP it produced significantly less than the amount it could generate using its preferred source ATP (Figs. 2 and 4). The ~35% drop in velocity, compared to ATP, could be due to either a decrease in myosin’s single molecule step size and/or a slowing of one or more of the kinetic steps that occurs while myosin is bound to actin(47). The results of the single molecule laser trap assay revealed slower kinetics and possibly a reduced step size (Fig. 4) explain the slower velocity. While the effect on steps size was small (7 vs. 9 nm for *ortho*-AzoTP vs. ATP), the effect on event lifetime was much more striking at 10 μM AzoTP (Figs. 2b and 2d). Indeed, even at a five-fold higher substrate concentration myosin stayed bound to actin significantly longer than with ATP at 10uM (Fig. 2d). This is indicative of a slowed rate of the substrate binding compared to myosin’s ATP-binding and/or a slower rate of dissociation of myosin from the actin filament (Figs. 2 and 4), suggesting that myosin has a weaker affinity for *ortho*-AzoTP than for ATP, which is consistent with the higher K_M_ observed when measuring V_actin_ as a function of substrate concentration (Fig. 3).

In contrast to the slowed rate of substrate binding, our kinetic modeling also suggested that the rate of release of the diphosphate form of *ortho*-AzoTP (AzoDP) is accelerated compared to the release of ADP (Fig. 3), which would be consistent with a reduced affinity for AzoDP when myosin is strongly-bound to actin. Thus the 35% reduction in V_actin_ at saturating substrate concentrations is likely due to a reduction in myosin’s step size combined with a slowed rate of AzoTP induced dissociation of actomyosin, and this occurs despite an acceleration of the AzoDP release rate. This is intriguing because this particular kinetic step is thought to limit the contraction velocity in skeletal muscle(53) and thus suggests that *ortho*-AzoTP shifts the step that limits velocity from being substrate release to substrate binding.

The decrease in myosin’s force-generating capacity was similar in magnitude (Fig. 4) to the decrease in V_actin_ (Fig. 1), at roughly 65% of the value observed with ATP. Force at the molecular level is dependent on the product of the force of a single actomyosin cross-bridge and its duty ratio (i.e. the portion of its cross-bridge cycle spent strongly-bound to the actin filament) (52). Therefore the observed 35% reduction in the peak force-generating capacity (Fig. 4) could be due to a decrease in one or both of these parameters. The decrease in the step size (Fig. 2) could be explained by a drop in the force per cross-bridge, however the dramatic decrease in the frequency of binding events, as well as the increase in time between events (Fig. 4f) suggests that myosin attaches to actin more slowly in the presence of *ortho*-AzoTP. This is evidenced not only by lower average force, but also by the reduced frequency of binding events and the absence of any high force events (> 1pN, see Fig. 4c). Myosin’s attachment rate is governed by how quickly it can hydrolyze the substrate and how quickly it can go from weakly to strongly interacting with actin(54). Thus, once *ortho*-AzoTP binds to myosin’s active site and causes dissociation from actin, it may have difficulty achieving the proper configuration to break the gamma-phosphate bond in AzoTP(48). This would cause myosin to remain dissociated from actin for a longer period of time than it would with ATP. Therefore, in comparison to myosin’s cross-bridge cycle with ATP, these data suggest that at saturating *ortho*-AzoTP concentrations the most pronounced effect is a slower rate of hydrolysis that greatly increases the time spent off of actin, along with a subtle drop in myosin’s step size (Fig.4c). The steps of the cross-bridge cycle thought to be altered based on these findings are represented in the model of the cross-bridge cycle shown in Fig. 5.

Thus, despite being able to use this synthetic energy source to power force and motion, myosin is not able to harness as much of the energy as it can with ATP. This is interesting because nearly all of the contacts between myosin and ATP occur between the gamma phosphate and Mg^++^(55), and should thus interact with the active site in a manner similar to ATP, and possesses nearly the same free energy because it comes from the breakage of the same gamma-phosphate bond. Therefore, the observed altered function suggests that the base portion of ATP must also play a role in proper binding of the substrate to the active site, as well as the efficiency of energy transduction. Indeed, simply rotating the position of the triphosphate moiety on the phenyl ring profoundly impacted myosin function (Fig. 1).

To directly examine the nature of the interactions between the substrate and myosin we performed molecular dynamics simulations in which the substrate was docked in the active site and the nature of the interactions between myosin’s ATP-binding site and each substrate were qualitatively determined. In general, these simulations show that the *ortho* version of AzoTP maintains a binding pose in the active site that is quite similar to ATP (Supplemental movie 6), providing potential insight into why it is a much better substrate for myosin than either the *para* or *meta* versions of AzoTP. Thus the 35% decrease in force and velocity observed with *ortho*-AzoTP may be due to the subtle variations in its interactions with myosin.

### Para-AzoTP is an uncompetitive inhibitor of ATP

In contrast to the relatively high efficiency of energy transduction observed using the *ortho* form of AzoTP, myosin did not move in presence of the *para* isomer of AzoTP (Fig. 1). Interestingly, when mixed with ATP in the motility assay the *para* variant of AzoTP inhibited myosin function in a dose-dependent manner (Fig. 1d), reducing velocity by 50% when both substrates were set to an equal concentration of 2mM. The inhibition was readily reversible (Supplemental video 5), however it was less pronounced at lower ATP concentrations (Supplemental Fig. 2) as would be expected with a competitive inhibition mechanism(56). Indeed, the data are most consistent with an uncompetitive or anti-competitive mechanism(56). This form of inhibition putatively operates through a mechanism in which the inhibitor recognizes and traps the enzyme substrate complex, creating a branch in the canonical pathway(57). There are previous reports of uncompetitive inhibitors of myosin, however they typically act to prevent actomyosin binding, and as a consequence, actin-induced acceleration of P_i_ release from the active site, trapping myosin in an unbound or weakly bound state(58). With *para*-AzoTP the actin filaments still move at a very slow velocity at saturating *para*-AzoTP concentrations, suggesting that it inhibits product release on actin (e.g. ADP-release) and/or slows ATP-induced dissociation from actin. This is an important finding because the ability to inhibit myosin is a very powerful tool in biology(58), thus this provides researchers with a new inhibitor that affords the ability to more precisely characterize the mechanisms of cellular processes driven by myosin. The reversibility of the compound also presents an advantage over other compounds such as blebbistatin which are typically non-reversible(59).

Our molecular dynamics simulations with this compound were initiated with *para*-AzoTP in the active site, which may be a rare event in the in vitro motility assay, but nonetheless show that the triphosphate moiety can fit into myosin’s ATP-binding site. However, the simulations also indicate that this phenyl ring makes strong contact with a tyrosine residue near the ATP-binding site and can lead to significant deformation of the loop (Supplemental movie 6). Thus, it is somewhat surprising that despite being a triphosphate compound it does not appear to competitively inhibit myosin. One possible mechanism of action to explain this discrepancy is that *para*-AzoTP recognizes when myosin has a nucleotide in the active site (ADP or ATP) and traps myosin in actin bound state, as uncompetitive inhibitor (see Fig. 5).

**Figure 5.**
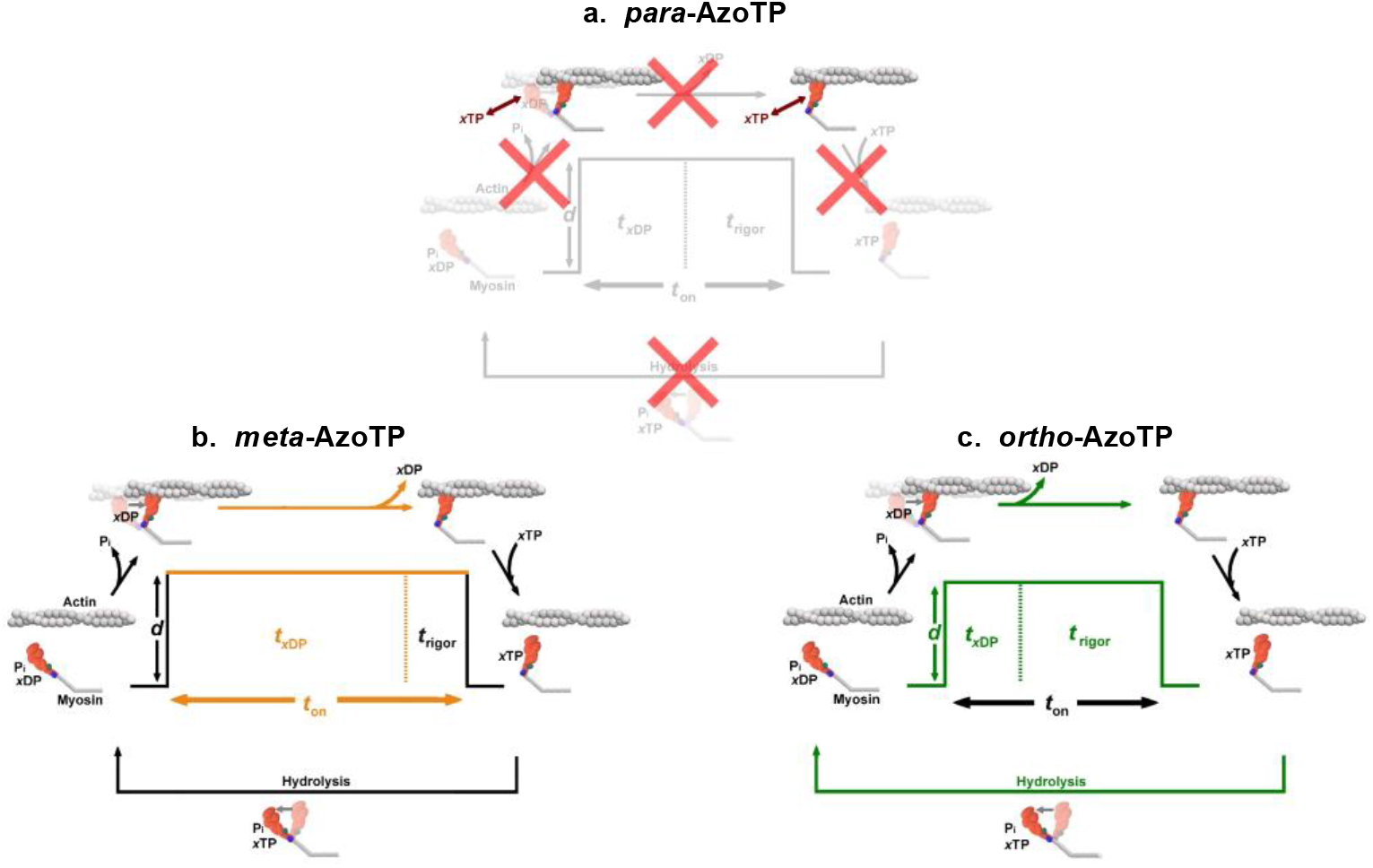
Hypothesized schemes of how each isomer affects myosin’s cross-bridge cycle. An equivalent model for ATP is shown in Figure 3. **a)** *para*-AzoTP traps filaments in a what appears to be a rigor-bound state (see Supplementary video 1) suggesting that *para*-AzoTP binds to actomyosin in the rigor state and prevents the release of the diphosphate form of the compound, or that *para*-AzoTP is does not bind to the active site. We favor the latter because the mechanism of inhibition appears to be uncompetitive (see Supplementary Fig. 5). The transitions marked with a red “X” are unlikely to occur. **b**) For *meta*-AzoTP the slowed velocity at saturating substrate concentrations (Fig. 1b) suggest that it was not due to a decrease in the percentage of moving filaments (Supplementary video 2 vs. 8) suggests that the diphosphate form of the substrate is released more slowly than ADP. This idea is depicted as a prolonged xDP lifetime. **c)** *Ortho*-AzoTP was used by myosin to generate force and motion but takes a smaller step (*d*) when binding to actin (Fig. 2c), has a prolonged rigor lifetime at low ATP (Fig. 2b), and at least one step off of actin is slowed because it generates ~35% less force than with ATP (Fig. 4).

### Meta-AzoTP greatly slows myosin’s velocity

Myosin’s mechanochemical cycle seems to be altered in a distinctly different way when *meta*-AzoTP is the substrate. Myosin moves actin but at only ~1% of the velocity of ATP, despite all the filaments moving in a continuous pattern (see Supplemental video 2), seemingly shifting it into a “low gear”. Velocity, in the motility assay, is believed to be limited by the rate at which myosin detaches from actin(51). This detachment rate is governed by how quickly myosin can release the diphosphate form of the substrate and then rebind a new triphosphate substrate. Therefore one, or both, of these parameters may be slowed with *meta*-AzoTP. However, the observation that the motion was continuous, combined with the observation that the V_actin_ is not increased by doubling the substrate concentration to 4mM (see Supplementary video 7 compared to Supplemental video 2) suggests that the slowed V_actin_ is not due to a prolonged rigor state. Rather, since velocity is thought to be limited by the rate of release of the diphosphate form of the substrate at saturating substrate concentrations(53) this suggests that myosin is trapped in a state waiting for the active site to release the *meta*-AzoDP (Fig. 5). Thus myosin might have a very high affinity for the *meta* form of AzoDP, which would imply that this fast skeletal muscle myosin could be manipulated to take on the properties of a much slower myosin, such as smooth muscle myosin(47), by simply changing the isomer of the substrate. Since the structural change to the substrate is relatively minor yet the effect on myosin is pronounced, control of the motor in this way could afford researchers with a powerful tool to gain precise detail of the mechanisms that govern, and gate, substrate binding and release from the active site. More broadly, it could be used to provide new insights into how myosin transduces chemical energy from the breakage of the gamma bond of a triphosphate-based compound into force and motion.

## Conclusion

Myosin, and evolutionarily-related ATPases, have evolved over billions of years to bind to, and harness the energy of, ATP hydrolysis to perform various intracellular tasks(60). Thus, it is not surprising that it has been quite challenging to develop an alternative form of energy that myosin can harness to transduce chemical energy into mechanical work as efficiently as ATP(5). Despite this challenge we have developed an abiotic, non-nucleoside triphosphate that muscle myosin can use to drive actin filament motion (Fig. 1) and generate mechanical work (Fig. 4). And while myosin generated less force and velocity than with ATP it is quite surprising, given the stark differences in the structure (Fig. 1), that the azobenzene triphosphate can power myosin’s powerstroke and cyclic interactions with an actin filament. Despite the reduced efficiency of the substrate (e.g. *ortho*-AzoTP vs. ATP) these findings present a powerful avenue to gain control over myosin’s function in two ways; first by affecting specific steps in myosin’s cross-bridge cycle, what we refer to as intrinsic control, and second by using the light-sensitive azobenzene molecule to control by light, what we consider to be extrinsic control. This is therefore important information for ongoing efforts to gain control over molecular motors both in a cellular setting and in synthetic nanodevices. This approach will also be a powerful method to gain novel insight into the fundamental rules that govern the energy transduction process in myosin and related molecular motors.

## Author contributions

M.W. performed research, analyzed data, and prepared the manuscript; E.O. synthesized the reagents, performed research and helped prepare the manuscript; S.P.J. designed and synthesized the reagents; X. L. performed research, B.S. performed research and analyzed data; M.U. performed research and analyzed data; J.C. designed research and helped write the manuscript; D.V. designed research, helped synthesize reagents and prepare the manuscript; E.P.D. designed research, performed research, analyzed data and prepared the manuscript.

## Acknowledgements

This work was support by an Innovative Project Award from the American Heart Association (Grant # 18IPA34170048) to E.P.D., D.V and J.C.

